# Highly synchronized inhibition from Purkinje cells entrains cerebellar output in zebrafish

**DOI:** 10.1101/2024.06.27.600928

**Authors:** Vandana Agarwal, Sriram Narayanan, Mohini Sengupta, Aalok Varma, Sudeepta Sarkar, Suma Chinta, Vatsala Thirumalai

## Abstract

Cerebellar function, known to be important for motor learning and motor coordination, is mediated by efferent neurons that project to diverse motor areas. To understand cerebellar function, it is imperative to study how these efferent neurons integrate inputs from the principal neurons of the cerebellar cortex, the inhibitory Purkinje neurons (PNs). In zebrafish, PNs are bistable and we show here that bistability influences spike synchrony among PNs. Bistability also alters spike correlation with motor bouts. We asked how PN population synchrony influences Eurydendroid cells (ECs), which are postsynaptic targets of PNs and are the cerebellar efferent cells in zebrafish. Using optogenetics, we artificially modulated population synchrony of PNs over millisecond time scales and showed that under conditions of high synchrony, EC firing is briefly suppressed and entrained by PN spiking. However, the magnitude of such modulation is relatively small and indicates a strong combined influence of other synaptic inputs on EC spiking.

**Key Points:**

- Cerebellar Purkinje neurons (PN) in larval zebrafish alter simple spike correlations with each other based on cellular state.
- They also alter simple spike correlations with motor bouts as a function of state.
- We altered PN population synchrony in a graded manner using optogenetics.
- PN targets are cerebellar efferent neurons, which in teleosts are called eurydendroid cells.
- When PN population is firing with high synchrony, eurydendroid cells are entrained better than when the PN input is asynchronous.
- This can explain how PNs use bistability to modulate their influence on cerebellar output and ultimately, motor behavior.

## Introduction

Cerebellar output is important for balance, planning and execution of movements, and in several non-motor functions as well (Thach, 1968; Koziol et al., 2014; Kunimatsu et al., 2018; Narayanan and Thirumalai, 2019; Kebschull et al., 2024). In mammals, this output is constituted by the projections of the deep cerebellar nuclei (DCN), while in teleosts, the homologous eurydendroid cells (ECs), undertake this function (Ikenaga et al., 2006; Bae et al., 2009; Heap et al., 2013). Yet, it is not clear how DCN/EC spike output is patterned or how they participate in cerebellar computation.

The cerebellum has well conserved circuitry from fish to mammals (Bell et al., 2008), underlining the importance of this circuit architecture for survival. Both in mammals and fish, Purkinje neurons (PNs) integrate a large number of weak excitatory inputs from granule cells (GCs) via the parallel fiber (PF) pathway and also few, but strong inputs from climbing fibers (CF) originating from the inferior olive (Nieuwenhuys, 1967). PNs make GABAergic, inhibitory projections to DCN/ECs such that these output cells receive convergent inhibition from many PNs, estimated to be 20 to 50 in mice (Person and Raman, 2012) and 1 to 3 in larval zebrafish (Harmon et al., 2020). While many features are conserved, there are also a few distinctions between mammalian and teleostean cerebella. In mammals, DCN receives direct inputs from mossy fibers and olivary climbing fibers (McCrea et al., 1977; Shinoda and Sugihara, 2013). In teleosts, ECs receive inputs from PFs of granule cells which are post-synaptic to mossy fibers and seem to not receive climbing fiber input (Ikenaga et al., 2006, 2023; Magnus et al., 2023). While in mammals, the DCN neurons are located ventrally within the cerebellar white matter, in teleosts such as zebrafish, the ECs are located proximal to the Purkinje cell layer, making them easily accessible for *in vivo* recordings and manipulation.

Several studies have explored the role of convergent PN input in modulating the spiking activity of cerebellar nuclear cells (McDevitt et al., 1987; Person and Raman, 2012; Sarnaik and Raman, 2018) as well as in regulating the kinematics of movements (Thach, 1968; Armstrong and Edgley, 1984a; McDevitt et al., 1987; Heck et al., 2007; Hoogland et al., 2015; Herzfeld et al., 2018; Sarnaik and Raman, 2018; Chang et al., 2020; Sedaghat-Nejad et al., 2022). However, given that PNs inhibit DCN/ECs and that both cell types are co-active during movement (Thach, 1968; Armstrong and Edgley, 1984a, 1984b), the influence of PNs on DCN/EC spiking is still nebulous. While some studies suggest that PN population synchrony entrains DCN spike timing and thus regulates motor kinematics (Heck et al., 2007; Person and Raman, 2012; Hoogland et al., 2015; Sarnaik and Raman, 2018; Sedaghat-Nejad et al., 2022), others propose that PNs communicate via a rate code (Payne et al., 2019; Herzfeld et al., 2023). We therefore sought to investigate the contribution of PN synchrony in modulating EC activity in vivo in larval zebrafish. Larval zebrafish are transparent and their ECs are dorsally located, thus allowing us to record from them while simultaneously manipulating PN activity using optogenetics.

Previously, we reported that PNs in larval zebrafish exhibit membrane potential bistability: they are naturally found at relatively depolarized or hyperpolarized membrane potentials and fire in distinct modes at these states. We found that PNs fire tonically when depolarized and in bursts when hyperpolarized (Sengupta and Thirumalai, 2015). We wished to explore how bistability affects the PN population and in turn how this shapes the cerebellar output. Here, we report that PN simple spikes are highly correlated with motor output only when they are in the bursting state. The same PN, when it is depolarized and is tonically spiking, loses the simple spike correlation with motor output. In addition, we find that simple spike correlations are higher between proximate PNs when they are both bursting compared to when one or both of them are tonic. This suggests that PNs could use bistability as a way of modulating population spike synchrony to engage with or disengage from motor output.

To explore the above idea further, we recorded from ECs during optogenetic stimulation of PNs. By varying the light stimulation intensity, we were able to modulate the probability of spiking in a graded fashion, which in turn translates to greater or lesser synchrony across the population. We find that when PNs are firing synchronously, ECs can be entrained to their firing while asynchronous firing of PNs is unable to affect EC spike times. Yet, we find that the effect of PN inhibition on ECs even under conditions of highest synchronicity is relatively modest, suggesting a significant role for non-PN inputs on cerebellar output.

## Materials and methods

### Animal care and use

All experiments were approved by the Institutional Animal Ethics Committee and the Institutional Biosafety Committee of the National Centre for Biological Sciences. Adult zebrafish were housed in a recirculating water system (ZebTEC, Tecniplast Inc). These adults were bred to obtain experimental fish that were used for experiments at 6-8dpf. Experimental fish were raised in either E3 medium (composition in mM: 5 NaCl, 0.17 KCl, 0.33 CaCl_2_, 0.33 MgSO_4_ and 0.5 Methylene blue; pH 7.2 to 7.4) or ZebTEC system water with 0.5mM methylene blue. They were kept in an incubator maintained at 28°C, with a 14:10h light:dark cycle.

### Generation of transgenic fish

For transient transgenesis, a mixture containing 25 ng/µL pTol2-Ca8-cfos:ChR2(H134R)-mCherry plasmid and 50 ng/µL Tol2 transposase mRNA (Urasaki et al., 2006) was microinjected in embryos from hspzGFFgDMC156A;UAS:GFP fish (received from Prof. Masahiko Hibi) (Takeuchi et al., 2015) at the 1- or 2-celled stage. Larvae were screened for GFP expression in ECs and sparse, mosaic expression of ChR2-mCherry in PNs.

To generate stable lines, a cocktail of 25 ng/µL plasmid containing pTol2-Ca8-cfos:ChR2(H134R)-mCherry construct flanked by Tol2 sequences and 50 ng/µL Tol2 transposase mRNA was microinjected into Indian wild type embryos at the 1- or 2-celled stage. These embryos were grown and screened for germline expression of ChR2-mCherry. Founder fish (F_0_) with germline expression were outcrossed with hspzGFFgDMC156A; UAS:GFP fish line to generate F_1_ fish that expressed ChR2-mCherry in PNs and GFP in ECs. F_1_ fish from different F_0_ parents were crossed to select for larvae with enhanced expression of ChR2-mCherry. After each breeding, fish with maximum expression of ChR2-mCherry were selectively raised and used for experiments. Only F_2_ and later generations of fish were used for experiments.

### Electrophysiology

To prepare larval zebrafish for electrical recordings, they were anesthetized using 0.01% Tricaine and pinned at the center of a Sylgard-lined recording chamber using fine tungsten wire pins (California Fine Wire). First two pins were inserted in the notochord – at the caudal end of the tail and near the swim bladder, third pin was inserted in the jaw and was used to orient the head dorsal up. Tricaine was washed off and replaced with external recording solution (composition in mM: 134 NaCl, 2.9 KCl, 1.2 MgCl_2_, 10 HEPES, 10 glucose, 2.1 CaCl_2_ and 0.01 D-tubocurarine; pH 7.8; 290 mOsm). 1 µM

TTX was added to the external solution for one experiment (as mentioned in the results section). The skin from the top of the head was removed to partially expose the brain and make the cerebellum accessible for patch clamp recordings from the PNs and ECs. Additionally, skin was also removed from the left side of the tail for experiments involving ventral root recordings.

Recording pipettes were pulled using P-97 and P-1000 micropipette pullers (Sutter Instruments). For patch recordings, filamented borosilicate capillaries with 1.5mm OD and 0.86mm ID (Warner Instruments) were used, whereas for ventral root recordings, non-filamented, thin-walled borosilicate capillaries with 1.5mm OD and 1.1mm ID (Sutter Instruments) were used. Patch and ventral root recording pipettes had tip resistances of 8-14MΩ and 1MΩ respectively. Pipettes were filled with filtered potassium gluconate based internal solution (composition in mM: 115 K gluconate, 15 KCl, 2 MgCl2, 10 HEPES, 10 EGTA, 4 MgATP; pH 7.2, 290 mOsm) for whole-cell recordings. For all the other recordings, filtered external solution was filled in the pipette.

The recording chamber was transferred to the electrophysiology rig and the sample was visualized using a 60x water immersion objective on an Olympus BX61WI microscope. Recordings were acquired using Multiclamp 700B amplifier, Digidata 1440A digitizer and pCLAMP software (Molecular Devices). Data was acquired at 20-50 kHz and low-pass filtered at 2 kHz. The amplifier gain was set to 1 for whole-cell recordings, whereas a gain of 20 to 1000 was used for extracellular recordings.

Whole-cell recordings were performed from PNs for examining motor bout correlations as a function of state and for verifying the functional expression of ChR2 in PNs. All the other recordings from PNs and ECs were performed in the loose-patch configuration and targeted using membrane ChR2-mCherry expression in PNs and cytoplasmic GFP in ECs for reference. For paired recordings, after obtaining loose-patch recording from one PN, the objective was lifted and a second electrode was brought into focus and maneuvered to the sample. A second cell close to the first recording was targeted for recording, trying to minimize tissue disruption. Once both channels produced stable signals, 1 minute and 10-minute-long recordings were obtained before a different cell was targeted for recording. Ventral root recordings (VRR) were performed from the tail of the fish and used as a proxy for swims. VRR were performed from the ipsilateral side along with simultaneous loose patch recordings from ECs.

### Optogenetic stimulation

Blue light was delivered using CoolLED pE-300^ultra^ for optogenetic stimulation. Light stimulation was provided either continuously or in pulses. For continuous light stimulation, the CoolLED pE-300^ultra^ was triggered by sending a TTL signal generated by Clampex (Molecular Devices). PNs were continuously stimulated with blue light either for 1s (Fig. 4C-top, D, E) or for 20s (Fig. 4C-bottom, F). For pulsed optogenetic stimulation, protocols were written using a graphical user interface that was written using Python. Each protocol contained multiple trials that were triggered by sending a TTL pulse using pCLAMP software. This trigger updated the trial information onto a Teensy 3.2 microcontroller, that turned the LED on and off as specified in the protocol. Pulses were 10 ms long and delivered at 20 Hz. These parameters were arrived at based on pilot experiments exploring a range of frequencies and durations. Pulsed stimulation during each trial lasted for either 1 or 3 seconds. The stimulation was performed at 6 different intensities, 0.16, 0.4, 0.7, 1.2, 3.0 and 4.6mW/mm^2^ for each cell, or as long as the cell could be stably patched. Light stimuli covered a region 290 µm wide, centered over the cerebellum.

### Data analysis

Spikes in PNs and ECs were detected in one of two ways: using template matching in Clampfit software or using custom Python scripts such as quickspikes (Meliza and Margoliash, 2012). Motor events were detected from the extracellular suction recordings by computing a rolling standard deviation of the recording with a 5ms window, followed by thresholding. Event detection was manually verified and the detection parameters adjusted manually if required. Events were aligned with optogenetic stimulation using TTL signals recorded parallelly on an analog input channel of the patch-clamp acquisition system. Further analysis of spike data was performed using custom analysis scripts written in Python and standard scientific computing libraries: *NumPy* (Harris et al., 2020), *SciPy* (Virtanen et al., 2020) and *statsmodels* (Seabold and Perktold, 2010). Visualizations were generated using matplotlib (Hunter, 2007) and Inkscape.

Motor signal-to-noise (motor SNR) was calculated by dividing the mean spike count from the start to 200ms after the initiation of motor bout by the median spike count of the cell and a threshold of 1.3 was used to classify a cell as motor tuned (Figure 1E). Linear regression was used to find a fit for the scatter (R^2^=0.62).

**Fig 1.**
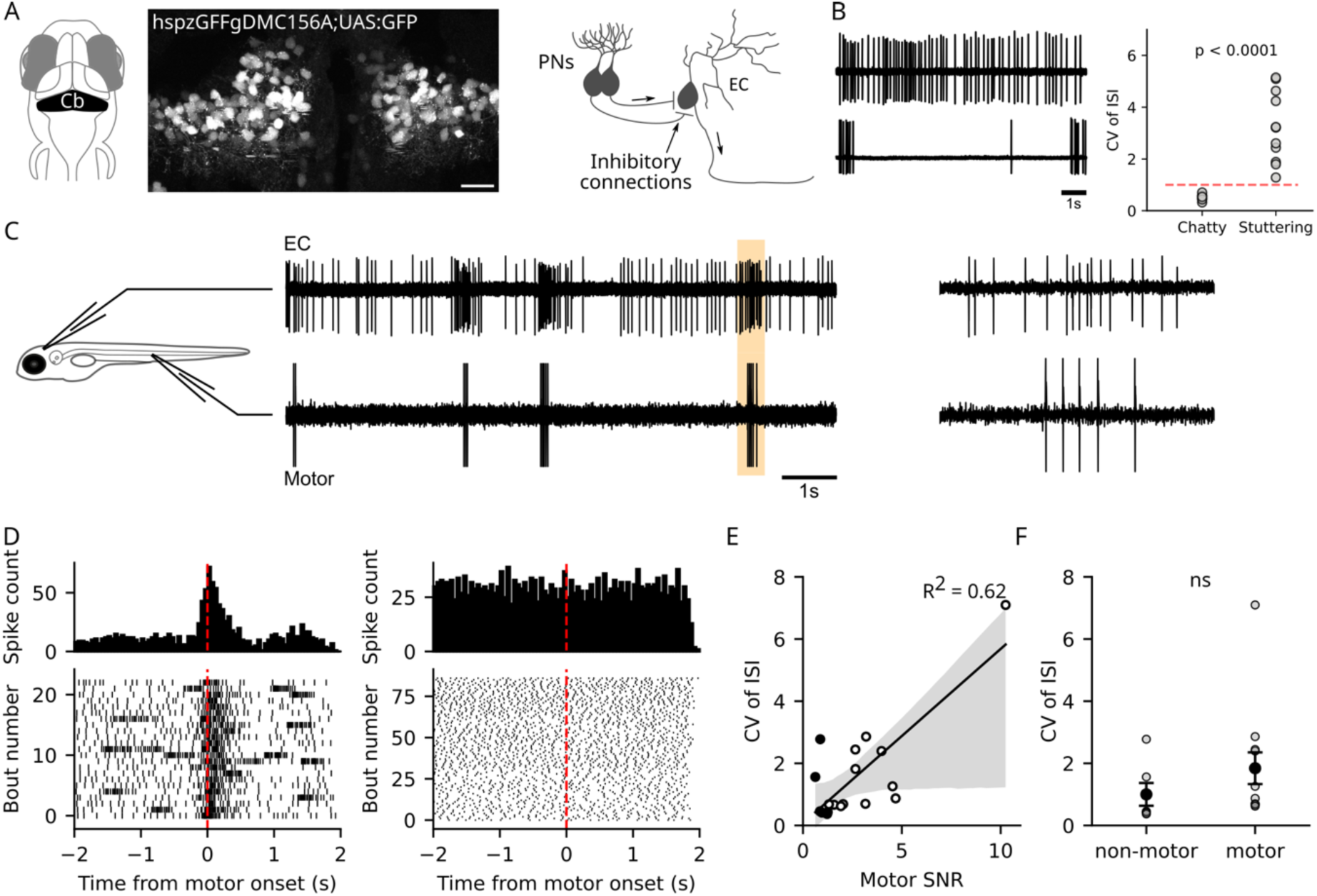
Modes of activation of ECs and their relationship with fictive motor output. A) Left: Schematic of fish head with cerebellum highlighted in black, Cb: cerebellum. Middle: Representative image showing GFP expression in ECs in larval zebrafish. Scale bar: 20µm. Right: Schematic depicting convergent inhibition onto ECs from PNs. B) Left: Representative recordings from chatty (top) and stuttering (bottom) ECs. Right: Distribution of the coefficient of variation of inter-spike intervals of chatty (n=8 cells) and stuttering ECs (n=11 cells) (p < 0.0001, Wilcoxon rank-sum test). Dashed red line: CV = 1, used as the decision boundary for classifying ECs as chatty or stuttering. C) Left: Representative simultaneous recordings from ECs in the cerebellum and suction recordings from tail. Right: Highlighted region of the recording on left (500ms) on an expanded scale. D) Left: Peri-event time histogram (top) and raster plots (bottom) of spikes in representative EC that showed modulation of activity with motor output (motor SNR=4.69). Same cell as in C. Right: Peri-event time histogram (top) and raster plots (bottom) for a representative EC that did not show changes in its activity with the motor bouts (motor SNR=0.96). Start of motor bouts has been aligned to zero. E) CV of ISIs of all cells versus motor signal-to-noise ratio. Black line: linear regression fit (R^2^=0.62), grey area: 95% confidence interval, filled circles: non-motor cells (n=6 cells), hollow circles: motor cells (n=12 cells). F) Distribution of CV of ISI of motor and non-motor cells. (ns:p-value=0.075, not significant, Wilcoxon rank-sum test).

Correlation coefficients between two PNs were calculated as follows. Instantaneous simple spike firing rate was calculated using the instantfr function in the MBL NSB Toolbox (https://github.com/wagenadl/mbl-nsb-toolbox), at a sampling frequency of 100Hz. The resulting output was then filtered using low-pass Butterworth filters with a time constant of 0.1s. Cross-correlations between two firing rate functions were calculated as Pearson’s product-moment correlation coefficient between the traces, with zero lag. To test whether the true cross-correlation could be obtained by chance, events from one of the two channels were scrambled and the resulting firing rate function computed. The correlation coefficient between this scrambled firing rate function and the true firing rate function of the other cell was then computed. This entire process was repeated 2000 times to obtain the distribution of “chance” correlation coefficients. The correlation was considered significant if the true correlation value lay outside the bounds of this “chance” distribution.

### Statistics

Statistical hypothesis testing was performed using the stats module in SciPy as well as the statsmodels Python package. The statistical tests used and their outcomes are presented in the respective figure legends.

## Results

### Eurydendroid cells are spontaneously active

We started with characterizing the baseline activity of ECs and their relationship with the motor output of the fish. Loose patch recordings were performed from GFP expressing ECs from 6-8 dpf hspzGFFgDMC156A; UAS:GFP larvae (Fig. 1A). Broadly we found two classes of ECs: cells that fired tonically, which we called ‘chatty’ and those that fired intermittently, which we called ‘stuttering’. To objectively classify cells as either chatty or stuttering, we calculated the coefficient of variation in their inter-spike intervals (CV^ISI^). While chatty cells had CV_ISI_ less than 1, stuttering cells had CV_ISI_ spread over a large range above 1 (Fig. 1B). Thus, we set CV_ISI_ = 1 as a boundary for clearly separating chatty and stuttering cells. Over a 5-minute recording period, we never observed chatty cells shift to stuttering activity or vice versa.

Previous studies have shown that ECs fire during spontaneous, evoked and conditioned motor bouts (Harmon et al., 2020; Najac et al., 2023). We wished to understand if chatty and stuttering cells exhibit distinct correlations with motor bouts. For this purpose, we recorded from ECs in the loose patch configuration while simultaneously recording spontaneous fictive motor bouts (Fig. 1C).

Majority of the ECs exhibited changes in their spiking activity around swims (12/18 cells), whereas a small fraction of cells did not show changes in activity during swims as compared to the baseline (6/18 cells). Motor-related ECs had bursts of spikes during swim bouts which is also evident from the peak in their peri-event time histogram and greater density of spikes in the raster plots around the start of swim bouts. No such modulation of spikes occurred for non-motor ECs (Fig. 1D). The coefficient of variation (CV) of ISI of motor tuned cells was greater than non-motor cells but the difference was not statistically significant (ns:p-value=0.075 not significant, Wilcoxon rank-sum test) (Fig. 1F). We therefore conclude that both chatty and stuttering ECs are likely to encode motor bouts in their spike patterns.

### Purkinje neuron state governs motor bout correlation and population synchrony

Our work has shown that larval zebrafish Purkinje neurons are also co-active during motor bouts and that these neurons are bistable: i.e., they can exist stably at one of two membrane potentials and therefore fire tonically or in bursts (Sengupta and Thirumalai, 2015). Purkinje neurons are inhibitory and therefore it is surprising that both Purkinje neurons and their post-synaptic targets, the ECs, are co-active during motor bouts. We hypothesized that the effect of Purkinje neuron inhibition could be to time lock EC spikes. Previous studies suggest that synchronous inhibition from Purkinje neurons is more effective than asynchronous inhibition at inhibiting DCNs, the mammalian homologs of ECs (Person and Raman, 2012). We propose that bistability in Purkinje neurons could be used to modulate Purkinje neuron synchrony and, therefore, to modulate EC spike times and correlations with the motor bouts. To test this, we first asked whether PN state determines motor bout correlations.

We recorded PN activity in whole cell mode while simultaneously recording the fictive motor bouts. The same cell was toggled to both the tonic and bursting mode through current injections and the incidence of various somatic events were monitored with respect to motor bouts. We found that simple spikes in PNs were highly correlated with motor bouts when in bursting mode (Fig. 2A, B). The same cell, when firing in tonic mode, lacked this correlation (Fig. 2C, D). CF inputs triggered all-or-none giant depolarizing potentials in the bursting state and broad calcium spikes in the tonic state. Both these events remained significantly correlated with motor bouts (Fig. 2A,B,C,D). These results suggest that PN state modulates simple spike correlations with motor bouts.

**Fig 2.**
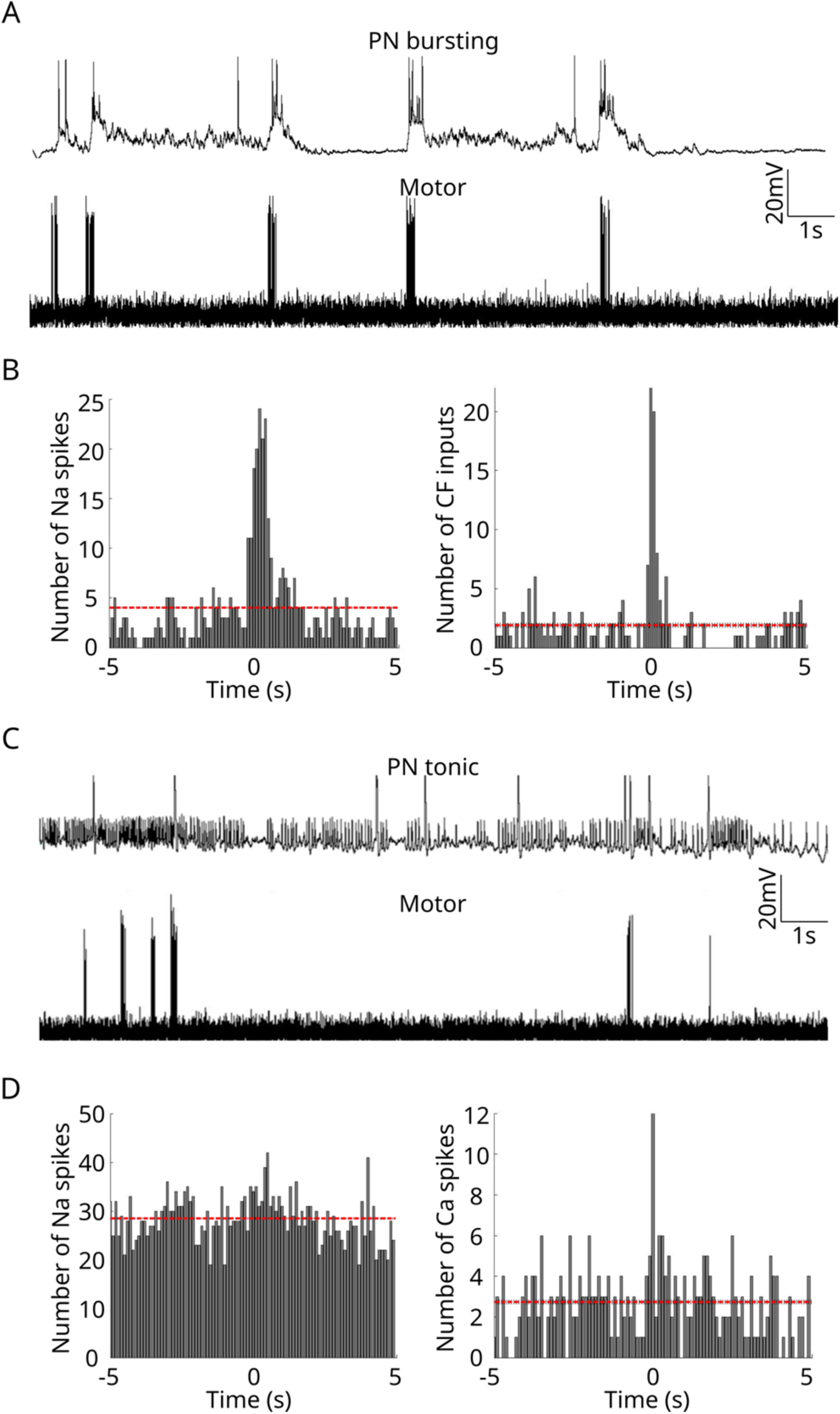
Activity of the PNs with respect to the start of motor bouts. A) Top: Representative current-clamp recording from bursting PN. Bottom: Simultaneously recorded fictive swims. B) Peri-event time histograms of sodium spikes (left), (p<0.01, chi-square test) and CF inputs (right), (p<0.01, chi-square test) of PNs in bursting mode with respect to the start of motor bouts (n=5 cells). C) Top: Representative current-clamp recording from the same PN as (A) showing tonic firing activity. Bottom: Simultaneously recorded fictive swims. D) Peri-event time histograms of sodium spikes (left), (ns: not significant, chi-square test) and calcium spikes (right), (p<0.01, chi-square test) of PNs in tonic mode with respect to the start of motor bouts (n=5 cells). Dashed line: mean, time zero on the peri-event time histogram represents the start of motor bouts.

Next, we wondered if bistability also affects population level synchrony among PNs, thereby affecting the strength of their influence on ECs. We made simultaneous loose patch recordings from pairs of proximal Purkinje neurons located within the same cerebellar hemisphere (Fig. 3A). Based on their simple spike activity, we classified them as bursting or tonic. Pairs of neurons were thus both bursting, both tonic or bursting and tonic (Fig. 3B, C, D). We first looked at the firing rate profiles of simple spikes from each neuron. We quantified the Pearson’s correlation coefficient between their firing rate profiles (Fig. 3E). From these data, it appears that the correlation in simple spike firing is highest in Bursting-Bursting pairs (median 0.5660) and lowest in Tonic-Tonic pairs (median 0.2912), with Tonic-Bursting pairs having intermediate correlations (median 0.3785). These values could not be obtained by chance after scrambling event times in one of the pair of PNs (Fig. 3F). These results suggest that cellular state of PNs does govern population synchrony.

**Fig 3.**
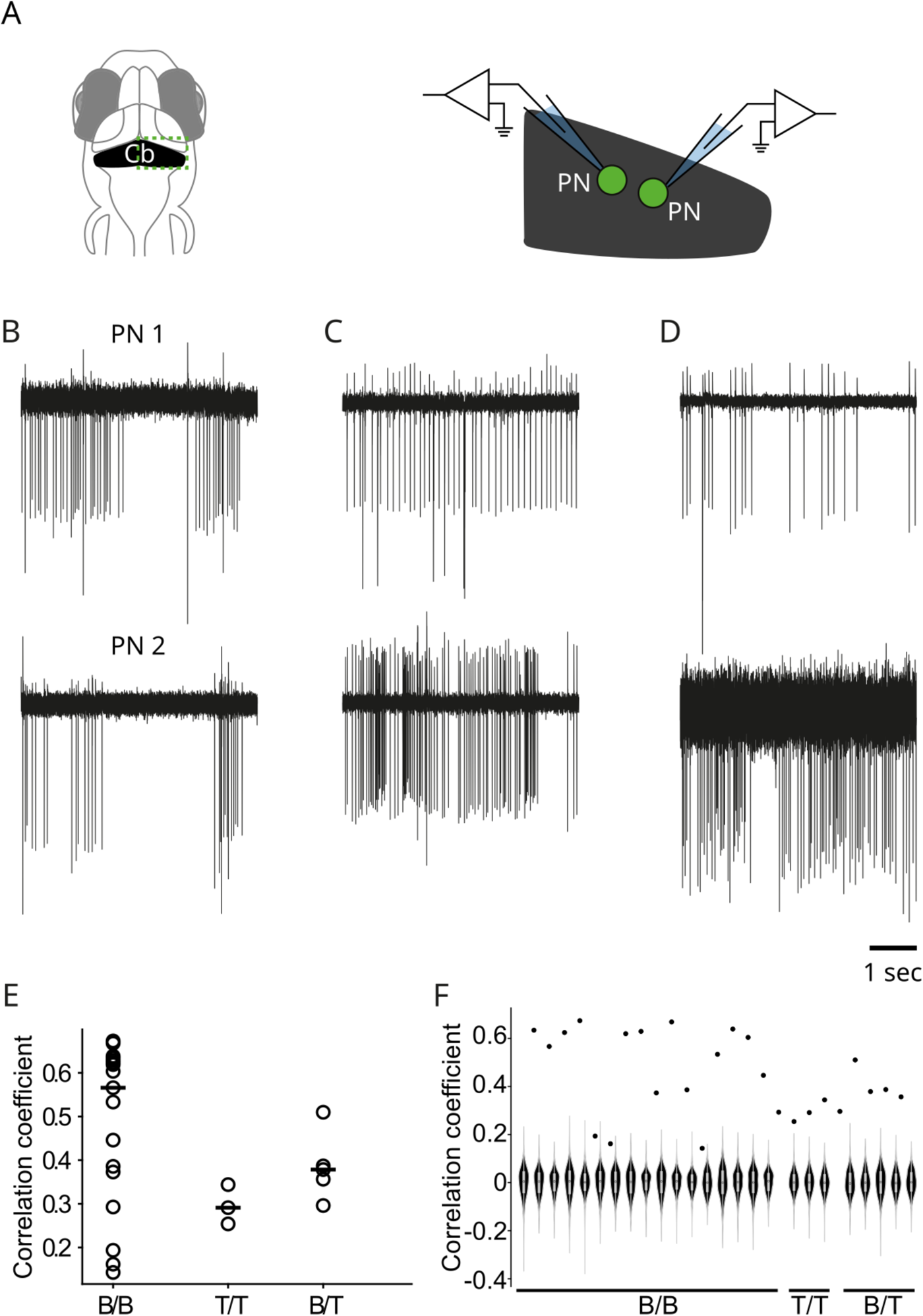
Simultaneously recorded PN pairs exhibit greater spike time synchrony during bursts. A) Left: Schematic showing larval zebrafish brain with cerebellum highlighted. Right: Schematic showing dual recording were performed from nearby PNs in the same cerebellar hemisphere. B) Representative recordings of a pair of bursting PNs recorded simultaneously. C) Same as B, for a pair of tonically active PNs. D) Same as B and C for a bursting-tonic pair of PNs. E) Pearson’s correlation coefficients between the firing rates for simple spikes for each kind of pair. F) True simple spike correlation coefficients (black circles) compared to the distribution of coefficients obtained by chance (violin plot), for each pair of neurons, sorted by their state combination. Correlations are significant if the true values lie outside the range of the violin plot. B and T are short for bursting and tonic respectively.

### Reliable and stable optogenetic excitation of PNs

Having established that simple spike correlations among PNs and between PNs and motor bouts are both functions of cellular state, we asked whether correlated or uncorrelated simple spike trains affect cerebellar output via ECs. We decided to manipulate population synchrony of simple spikes in Purkinje neurons to test effects on the cerebellar circuit. First, we needed to validate that we can specifically, reliably and stably excite PNs in larval zebrafish with milliseconds precision. To do so, we generated transient transgenic zebrafish in which ChR2(H134R), a channelrhodopsin variant (Berndt et al., 2011), was expressed in PNs under the Ca8 enhancer (Matsui et al., 2014). The ChR2(H134R) was fused with mCherry for ease of visualization (Fig. 4A). We screened larvae for mCherry expression and performed whole-cell patch clamp recordings from both ChR2-mCherry positive and negative Purkinje neurons (Fig. 4B, C). In the presence of tetrodotoxin (TTX), which blocks sodium channels and therefore all network drive, we found that 1s-long blue light pulses caused stable depolarization in ChR2 expressing PNs but not in ChR2-negative neurons (Fig. 4C, D). After cessation of blue light stimulation, ChR2-positive neurons returned to their pre-stimulation baseline. Interestingly, the quantum of depolarization during light pulse stimulation was proportional to the baseline membrane potential in these cells: more hyperpolarized the baseline membrane potential, greater the depolarization caused due to light stimulation and *vice versa* (Fig. 4E). These results show that PNs can be depolarized stably and reliably with optogenetic stimulation.

To test if the depolarization caused by ChR2 will be sufficient to induce spiking, we repeated the above experiment, but this time without TTX. We found that ChR2-expressing PNs do generate simple spikes when they are stimulated with blue light (Fig. 4C). The depolarization caused by stimulation was proportional to the baseline membrane potential in the spiking state as well. This meant that light stimulation brought PNs to near about -40mV, regardless of their baseline membrane potential (Fig. 4F).

### Optogenetic modulation of spike synchrony in PNs

After thus confirming that we can elicit reliable depolarization and spiking in larval zebrafish Purkinje neurons *in vivo*, we proceeded to develop a protocol to trigger spikes with variable probabilities, and therefore control spike time synchrony across the PN population. In this protocol, we used a train of brief pulses of light to control the time of PN spikes and varied the light intensities to modulate the effectiveness of stimulation. We hypothesized that higher light intensities will evoke more reliable responses from PNs and there will be an action potential every time a pulse of light is incident on the cerebellum, causing them to fire synchronously with millisecond time-scale precision. Whereas as we decrease the light intensity, the probability to elicit a spike in PNs will decrease. This will lead to greater variability in PN spike responses and times across the entire population, leading to their asynchronous activation.

We recorded responses of PNs to pulsed light stimulation for a range of light intensities namely, 0.16, 0.4, 0.7, 1.2, 3.0 and 4.6mW/mm^2^ (Fig. 5A), presented in a random order. We found that the average baseline firing rates as well were similar across all stimulation conditions (p = 0.829, Friedman test). Similarly, the average firing rate during the stimulation was also not different across the various light intensity conditions (p = 0.067, Friedman test) (Fig. 5B). However, at the level of single light pulses, at the strongest light intensity used, spikes were almost always elicited (mean±SEM: 0.85 ±0.04 spikes per pulse), while at the weakest intensity used, only a few trials were successful at eliciting a spike (0.155 ± 0.03 spikes per pulse). To be certain, we compared the mean firing rate before (pre-), during (in-) and after (post-) light pulses at every intensity and found that the stronger light intensities caused a significant increase in firing rate during the pulses compared to before or after (Fig. 5C).

**Fig 4.**
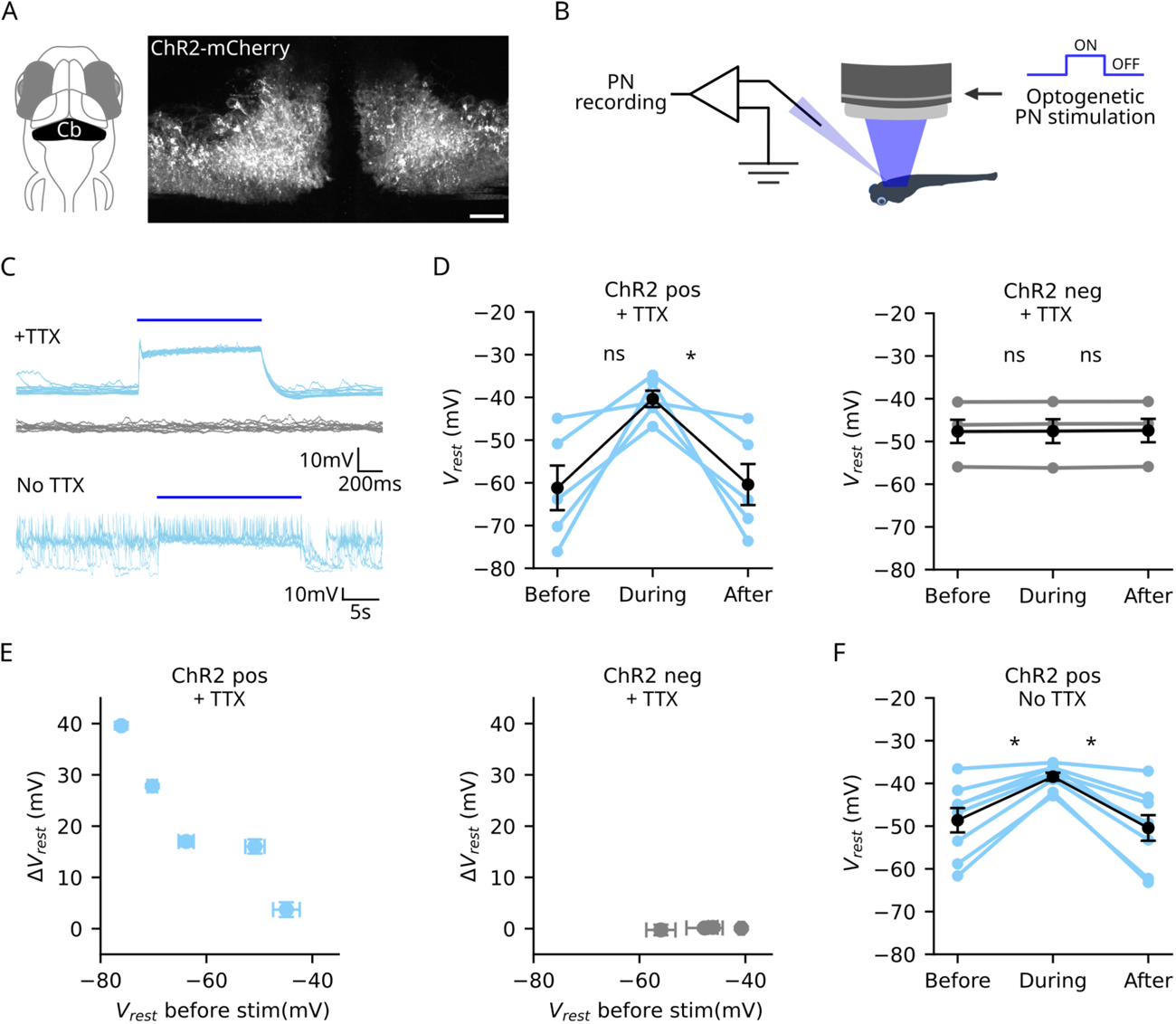
Optogenetic stimulation activates ChR2 expressing PNs. A) Left: Schematic of larval zebrafish head with the cerebellum highlighted in black, Cb: cerebellum. Right: Representative image showing ChR2-mCherry expression in the cerebellum. Scale bar: 20µm. B) Schematic showing the PN recording and optogenetic stimulation setup. C) Representative recordings from PNs. Top: whole cell voltage clamp recordings from ChR2 positive (blue traces) and ChR2 negative (grey traces) PN from same larva in the presence of TTX in the bath. 10 trials overlaid. Bottom: 5 trials from a ChR2 positive cell with no TTX in the bath. Blue bar above the traces represents the duration of light stimulation. D) Resting membrane potential of cells before, during and after blue light illumination in the presence of TTX. Left panel: n=5 ChR2 positive cells, right panel: n=4 ChR2 negative cells. *: p-value<0.05, ns:p-value>0.05, not significant, Friedman test. E) Difference in resting membrane potential between the stimulation ON period and the baseline resting membrane potential prior to stimulation onset, plotted as a function of the baseline value. F) Resting membrane potentials of cells before, during and after light stimulation in the absence of TTX. n=8 ChR2 positive cells. *: p-value<0.05, Friedman test. Blue: ChR2 positive cells, grey: ChR2 negative cells, black: averages of all the cells.

**Fig 5.**
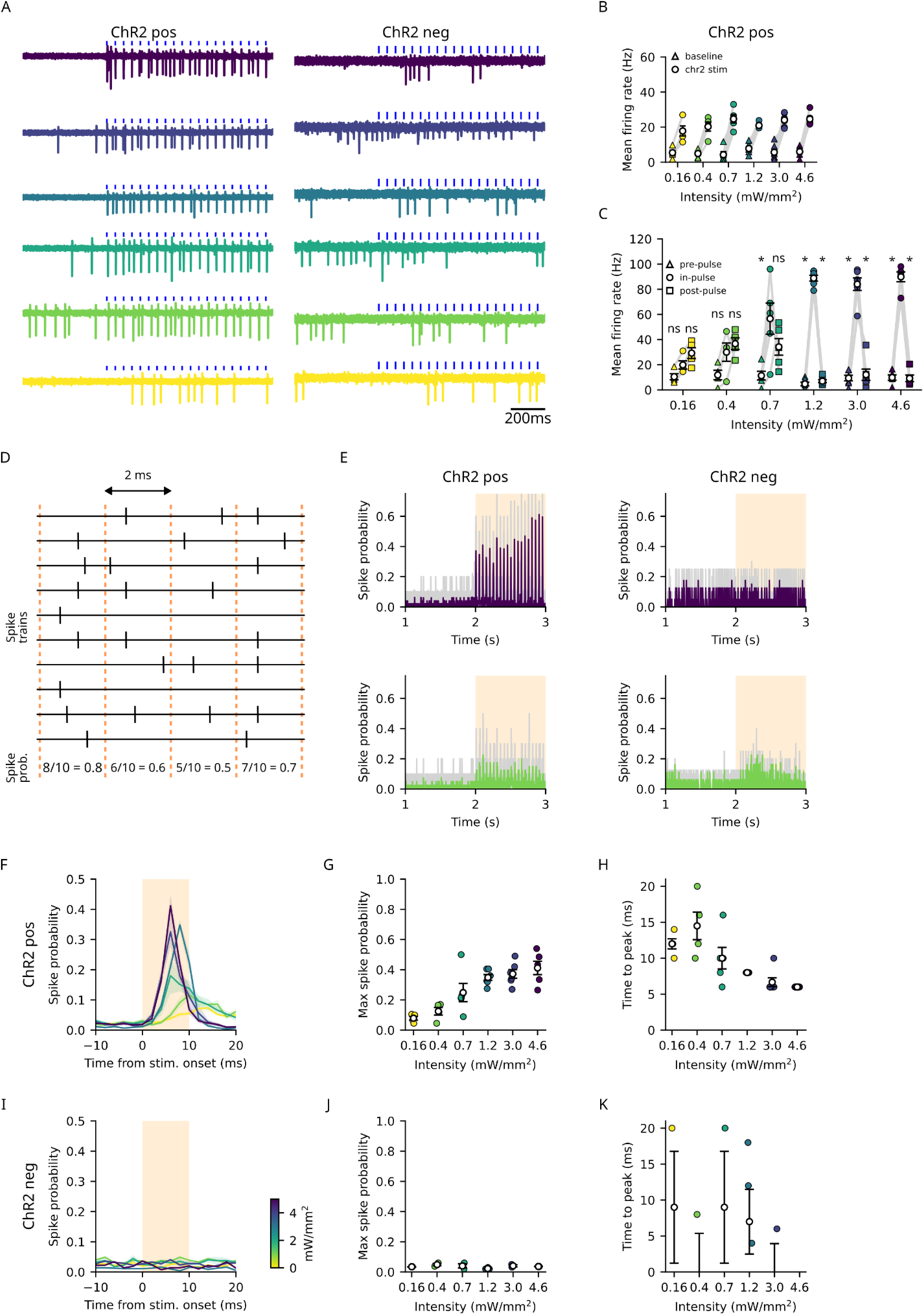
PN spikes are time-locked to pulsed optogenetic stimulation in an intensity dependent manner. A) Representative recordings from a ChR2 positive (left) and a negative PN (right) in response to pulsed light stimulation of varying intensities. B) Average firing rates of ChR2 positive cells before the start of optogenetic stimulation (triangles) and during stimulation (filled circles) for all the light intensities used. Firing rates before stimulation for all intensities, p-value=0.829, firing rates during stimulation for all intensities, p-value=0.067, Friedman test. C) Average firing rates of cells within 10ms windows before, during and after each pulse of light for ChR2 positive cells. *: p-value<0.05, ns:p-value>0.05, not significant, repeated measures ANOVA, Tukey’s post-hoc test. D) Schematic describing spike probability calculation. E) Plots showing average spike probability for all the cells at 4.6mW/mm^2^ (top) and 0.4mW/mm^2^ (bottom) light intensities for ChR2 positive (left panels) and ChR2 negative (right panels) cells. Grey traces: Spike probabilities of individual cells, coloured traces: Averages for all the cells. Start of optogenetic stimulation is marked as 2s. F) Pulse triggered spike probabilities of ChR2 positive cells for different stimulation intensities. Coloured traces: averages of all cells, shaded region around traces: SEM, shaded rectangle: duration of pulse. G) Peak value of spike probability during pulse, p-value=2.109e-5, GLM and H) time taken to reach maximum spike probability from the start of the pulse, p-value=1.120e-5, GLM for each intensity for ChR2 positive cells. I, J and K) same as F, G and H for ChR2 negative cells. J) p-value=0.961, GLM, K) p-value=0.921, GLM. Coloured symbols: cell averages, open black circles: averages of all cells, error bars: SEM, colour bar: intensities of light used for optogenetic stimulation. n=8 (4,4,5,6,6,5) and 7 (2,2,2,4,2,2) cells from ChR2 positive and negative larvae respectively for min to max stimulation intensities used.

From the above, it is clear that optogenetic stimulation of ChR2-positive neurons leads to spiking, a result which is important to establish, but not surprising. However, to assay whether optogenetic stimulation affects spike synchrony, it is important to test if the spikes are time-locked within the duration of each pulse. We calculated spike probability in 2 ms bins for the entire duration of the recording (Fig. 5D). For the maximum light intensities, an elevation in spike probabilities was observed in response to each pulse of light. This modulation was weaker at lower intensities. ChR2-negative cells showed no appreciable change in spike probability even at the highest light intensity used (Fig. 5E). Averaging across pulses, trials and cells, we noticed that the maximum spike probability was significantly higher for the highest light intensity compared to the lower intensities (Fig. 5F, G). The time to reach maximum spike probability was shorter for the highest light intensity and higher for the lower intensities (Fig. 5H). Such modulation in firing rate was not present for any of the light intensities used in ChR2-negative cells (Fig. 5I-K).

These experiments demonstrated that by modulating the intensity of blue light, we can dial up or down the probability of spiking and the latency to spike in ChR2-positive Purkinje neurons. At the strongest light intensity of 4.6 mW/mm^2^, we could elicit spikes that are time-locked in millisecond time scale, reliably. Across the different light intensities used, the mean firing rate remained similar, however, the time locking of spikes was different, thus allowing us to probe the effect of spike synchrony without the confounding effects of spike rates.

### Synchronous PN activity induces phase shifted modulation of spiking in ECs

After developing the protocol to optogenetically modulate population synchrony of PNs, we studied the effect of convergent synchronous PN inhibition onto single ECs. We recorded activity of ECs using loose patch recordings while simultaneously activating PN population. No evident difference in the spiking activity of ECs was observed in raw activity traces in response to different levels of PN synchrony (Fig. 6A). Average firing rates of ECs for the entire duration of pulsed stimulation also did not show consistent change for the stimulation protocols used (Fig. 6B). Number of spikes per pulse of light in ECs did not change significantly with intensity (Fig. 6C). Average spike probability traces of ECs for different light intensities also did not show clear modulation, unlike PNs (Fig. 6D). However, when we plotted pulse triggered spike probabilities for ECs, we observed clear modulation of EC spike probabilities as a function of light intensity. In ChR2 positive animals, ECs showed a dip in spike probability immediately at the end of the light pulse, and this was absent in ChR2 negative animals (Fig. 6E, F). We computed maximum change in pulse-triggered spike probabilities of ECs by subtracting the peak and trough values of the spike probability curves. ECs from ChR2 positive animals exhibited significant changes in spike probabilities as a function of the light intensity. Spike rate modulation seen with weak light intensity pulses was similar to chance level modulation in ChR2 negative larvae and was not statistically significant (Fig. 6G). Thus, ECs show significant modulation of spike probability in response to synchronous PN stimulation and this modulation occurred in the milliseconds timescale.

**Fig 6.**
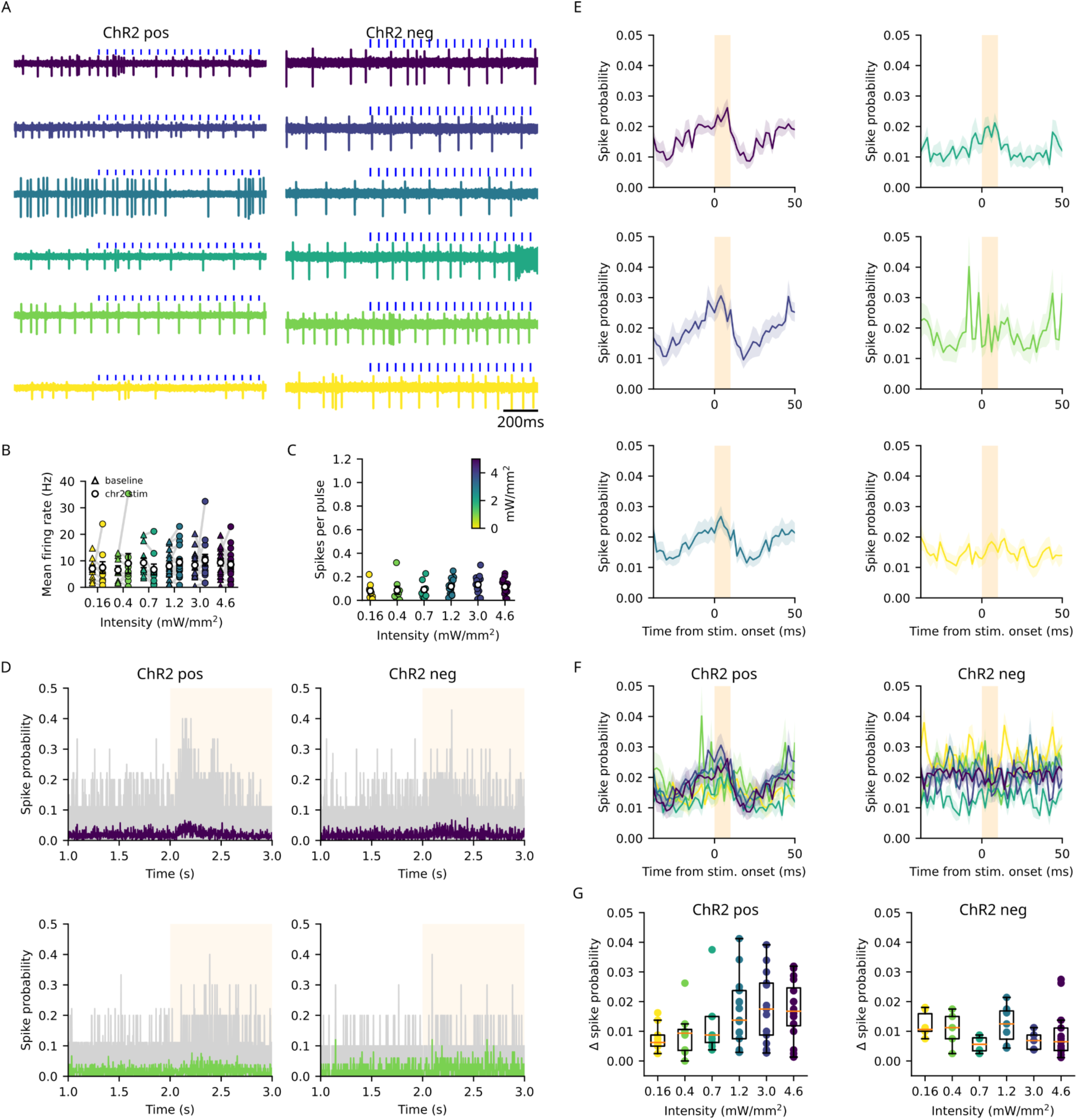
ECs show phase-shifted modulation of spiking activity in response to PN stimulation. A) Representative recordings of ECs from larvae expressing ChR2 in PNs (left) and with no ChR2 expression (right). Blue dashes above the traces indicate light stimulation. B) Plot of baseline mean firing rate of ECs before stimulation and average firing rate during stimulation from ChR2 positive larvae. C) Average number of spikes in ECs during pulses of light for different intensities of illumination. p-value=0.183, GLM D) Spike probability of ECs at maximum (4.6mW/mm^2^) (top) and 0.4mW/mm^2^ (bottom) intensity in ChR2 positive (left) and negative (right) larvae. Grey traces: spike probabilities of individual cells. E) Pulse triggered spike probabilities of ECs with varying levels of PN synchrony from ChR2 positive animals. 0 marks the start of the pulse and shaded rectangle indicates duration of stimulation pulse. Shaded region around traces: SEM. D, E and F) Coloured traces: population average. F) Same data as in E, pulse triggered spike probabilities at different light intensities, overlaid in ChR2 positive (left) and ChR2 negative (right) larvae. G) Maximum change in pulse triggered spike probabilities of ECs for all light intensities for ChR2 positive (left, p-value=0.012, GLM) and negative (right, p-value=0.443, GLM) animals. Coloured symbols: cell averages, open circles: population averages, error bars: SEM, colour bar: intensities of light used for optogenetic stimulation. n=23 and 16 cells from ChR2 positive and negative larvae respectively.

## Discussion

In this manuscript we have shown that in larval zebrafish, PN population synchrony and their spike correlations with motor bouts are a function of the cellular state. Further, synchronous firing of PNs over milliseconds timescales can entrain EC input by providing a short window of inhibition while asynchronous PN spiking has little effect on EC output. These results suggest that PNs in larval zebrafish could employ modulation of cellular state to alter their influence on ECs and thereby alter their relationship with motor output. Nevertheless, the extent of spike rate modulation in ECs by synchronous PNs was relatively modest thus strongly suggesting significant roles for other synaptic inputs.

### ECs share many properties with mammalian DCNs

We could distinguish two classes of ECs based on their spontaneous firing behavior: chatty and stuttering. We also find that a majority of ECs we sampled showed an increase in spike rates correlated with the start of motor bouts irrespective of whether they were chatty or stuttering. We believe these are distinct classes of ECs and not different firing modes of the same neurons since we never observed a switch in the firing mode even during long duration recordings. As we were recording in loose-patch rather than whole-cell mode, we were not dialyzing the cellular contents and therefore, these are most likely the natural activity patterns intrinsic to these cells. A previous study reported silent and intrinsically spiking ECs using whole cell recordings. The silent cells had depolarized membrane potentials compared to the spiking cells (Harmon et al., 2020). Further investigation is necessary to delineate the prevalence, distribution and mechanistic origins of these EC cell types.

At least two molecularly distinct subpopulations of ECs have been identified: olig2^+^ and calb2b^+^ ECs, each occupying distinct regions in the dorsoventral and mediolateral axes of the cerebellum, with the 156A gal4 line that we used in this study, labeling both olig2^+^ and olig2^-^ populations (McFarland et al., 2008; Bae et al., 2009; Takeuchi et al., 2015). Zebrafish ECs have also been shown to have diverse dendritic morphology and projection patterns and the projection targets of ECs correlate with their location within the cerebellum, suggesting a molecular code for the topographic organization of different EC types (Heap et al., 2013; Magnus et al., 2023). We do not yet know if the chatty and stuttering types we observe correspond to these molecularly and topographically distinct populations of ECs but such heterogeneity in EC populations will not be surprising.

It is noteworthy that ECs in ray-finned fish such as zebrafish, are homologs of the mammalian DCNs, yet, these cells are not organized into nuclei (Finger, 1978; Bae et al., 2009). Cartilaginous fish and amphibians have a single nucleus in each cerebellar hemisphere, while reptiles and birds have two, and mammals, three (Kebschull et al., 2020; Yopak et al., 2020). Single nucleus RNAseq analysis of DCN neurons in birds and mammals suggests that all cell types in these nuclei have been derived via duplication and divergence of two major classes of excitatory cells and three classes of inhibitory cells (Kebschull et al., 2020).

DCNs are known to be heterogenous with respect to their projection patterns, molecular expression patterns, firing properties and transmitter phenotypes. While DCNs consist of glutamatergic, GABAergic and glycinergic neurons (Bagnall et al., 2009; Uusisaari and Knöpfel, 2012; Kebschull et al., 2024), no inhibitory subtypes have been identified among ECs in teleosts and they seem to be a fully glutamatergic, long range projecting neuronal class (Bae et al., 2009; Heap et al., 2013; Matsui et al., 2014; Takeuchi et al., 2015; Harmon et al., 2020; Magnus et al., 2023). Given the above, one might speculate that distinct EC populations in fish might be related to ancestral glutamatergic populations observed in amniotes.

### PN versus non-PN influences on DCN/EC spiking

DCN/ECs are spontaneously firing cells and receive high frequency excitatory and inhibitory inputs. While mammalian DCNs receive mossy fiber and climbing fiber excitatory inputs (McCrea et al., 1977; Shinoda and Sugihara, 2013), fish ECs receive granule cell excitatory inputs (Ikenaga et al., 2006, 2023; Magnus et al., 2023). Also, while mammalian DCNs receive local GABAergic inhibition from other DCNs, in addition to PN inhibition, fish ECs appear to receive only PN inhibition (Bae et al., 2009; Heap et al., 2013; Kebschull et al., 2020, 2024). With their dendrites in the molecular layer, it is possible that they receive inhibition from molecular layer interneurons. This remains to be verified. ECs receive high frequency excitatory (EPSC) and inhibitory (IPSC) synaptic currents during spontaneous and evoked swimming. Interestingly, both EPSCs and IPSCs start to increase before the onset of spontaneous swimming but with a lag after the onset of evoked swimming (Harmon et al., 2020). This indicates that such synaptic currents convey signals about motor planning to ECs when spontaneous swimming bouts are about to occur. Further, while both simple spikes and CF inputs contribute to IPSCs in ECs, inhibition resulting from simple spikes is far more dominant (Harmon et al., 2020). This implies that synchronization of simple spikes among PNs could trigger strong inhibition of ECs. Lastly, both mammalian DCNs and ECs of adult zebrafish exhibit rebound spiking (Jahnsen, 1986; Aizenman and Linden, 1999; Magnus et al., 2023), suggesting that strong, synchronized inhibition could trigger timed spikes in these cells.

We find that synchronous inhibition from PNs generates a brief window of strong inhibition of spiking in ECs. ECs had a mean firing rate of 8.25 ± 0.6 (mean±SEM) Hz before optogenetic stimulation, similar to spontaneous firing rates of 7.9 ± 1.5 Hz reported by Harmon et al., 2020. Upon optogenetic stimulation of PNs with brief pulses, EC spike probabilities decreased by more than two-fold from 0.009 ± 0.003 to 0.026 ± 0.003. This decrease was present only for the strongest light intensities which cause synchronous firing in the PNs and not for the weaker intensities tested. However, given the relatively low rate of spiking in these cells, the absolute spike probability during this time window is small. Whether these modest increases in spike probability translate to meaningful behavioral outcomes remains to be seen.

### PN and EC contributions to motor planning and error correction

Here, we have shown that both PNs and ECs in larval zebrafish are co-active during motor bouts. This has been observed by our previous studies and other investigators as well (Sengupta and Thirumalai, 2015; Scalise et al., 2016; Harmon et al., 2017, 2020; Knogler et al., 2019; Najac et al., 2023). Our optogenetic stimulation experiments show that PNs can modulate EC spiking at the tens of milliseconds timescale, thus sculpting EC output precisely during ongoing motor bouts, which typically last over several hundred milliseconds to seconds. The precise computation performed by the cerebellar circuit remains to be solved. Yet, a few clues have been revealed. First, in larval zebrafish, PN correlations with motor bouts is a function of cellular state. Though existence of bistability of PNs is highly debated in mammals (Loewenstein et al., 2005; Schonewille et al., 2006; Engbers et al., 2013), it is clearly present in larval zebrafish (Sengupta and Thirumalai, 2015; Knogler et al., 2019). There are far fewer PNs in zebrafish compared to mammals and bistability could be a feature of PNs in teleosts to increase computational flexibility with limited neuronal hardware. Nevertheless, it offers us an opportunity to address how neurons might use bistability to perform complex computations. In this context, it appears that larval zebrafish PNs switch states to opt in or out of participating in motor tasks. Even in such cases, only the correlations with simple spikes are altered, while leaving CF input correlations relatively intact.

What distinct information is conveyed by the simple spikes versus the CF inputs? We found that when larval zebrafish are presented repeating visual stimuli, expectation for that stimulus is encoded by both simple spikes and CF inputs. Both these signals feed into motor planning circuits leading to shorter latency to generate motor bouts. When an unexpected stimulus occurs, the error in the expectation generates a much larger signal in PNs, possibly by the CF inputs. This error is then used to update expectation based on the current stimulus (Narayanan et al., 2024). These processes would depend on the simple spikes and CF inputs generating timed IPSCs in ECs via synchronization of the PN population, as demonstrated in the current study.

These results are in agreement with studies on mammalian DCNs. In mice, optogenetic manipulation of PNs that interferes with normal spike rate modulation of DCNs results in increased locomotor ‘slips’, while the same manipulations delivered such that the normal spike rate modulation was allowed, caused fewer such slips (Sarnaik and Raman, 2018). Thus, the precise temporal modulation of DCN/EC spikes by synchronized PN inhibition is key to planning and controlling motor output (Narayanan and Thirumalai, 2019). Future studies will have to delineate how DCN/EC output feeds into the sensory expectation and motor planning circuits.

